# Screening for Potential Interaction Partners with Surface Plasmon Resonance Imaging Coupled to MALDI Mass Spectrometry

**DOI:** 10.1101/2020.05.07.082776

**Authors:** Ulrike Anders, Maya Gulotti-Georgieva, Susann Zelger-Paulus, Fatima-Ezzahra Hibti, Chiraz Frydman, Detlev Suckau, Roland K.O. Sigel, Renato Zenobi

## Abstract

We coupled SPR imaging (SPRi) with matrix-assisted laser desorption/ionization mass spectrometry (MALDI MS) to identify new potential RNA binders. Here, we improve this powerful method, especially by optimizing the proteolytic digestion (type of reducing agent, its concentration, and incubation time), to work with complex mixtures, specifically a lysate of the rough mitochondrial fraction from yeast. The advantages of this hyphenated method compared to column-based or separate analyses are (i) rapid and direct visual readout from the SPRi array, (ii) possibility of high-throughput analysis of different interactions in parallel, (iii) high sensitivity, and (iv) no sample loss or contamination due to elution or micro-recovery procedures. The model system used is a catalytically active RNA (group IIB intron from *Saccharomyces cerevisiae, Sc*.ai5γ) and its cofactor Mss116. The protein supports the RNA folding process and thereby the subsequent excision of the intronic RNA from the coding part. Using the novel approach of coupling SPR with MALDI MS, we report the identification of potential RNA-binding proteins from a crude yeast mitochondrial lysate in a non-targeted approach. Our results show that proteins other than the well-known cofactor Mss116 interact with *Sc*.ai5γ (Dbp8, Prp8, Mrp13, and Cullin-3), suggesting that the intron folding and splicing are regulated by more than one cofactor *in vivo*.

**Graphical abstract:** 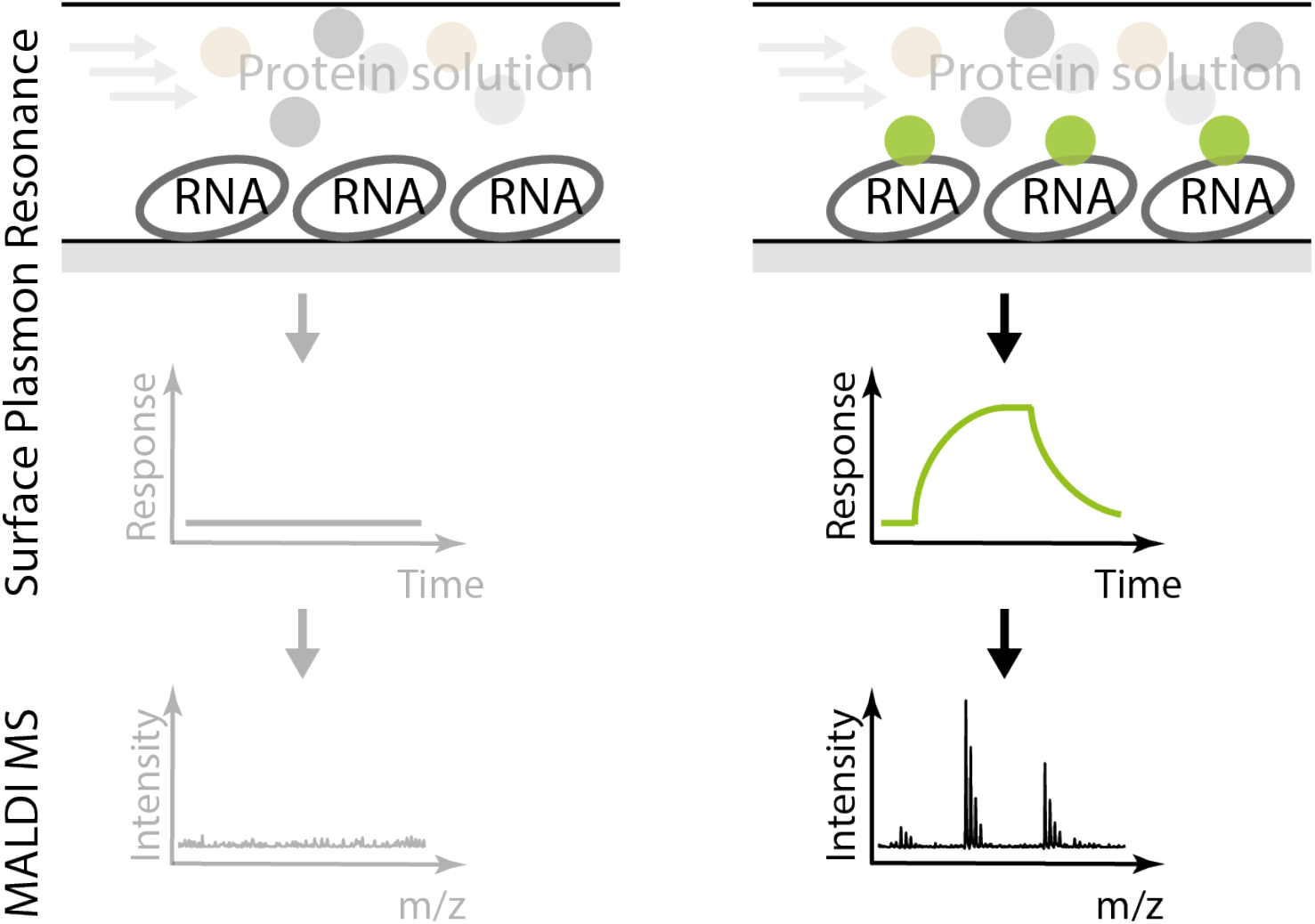

**Author contribution:** Ulrike Anders and Maya Gulotti-Georgieva contributed equally to this work.

## 1. Introduction

To identify molecular partners and simultaneously determine the binding affinity, a combination of several analytical methods would usually be required, rendering work laborious and time-consuming. The coupling of surface plasmon resonance imaging (SPRi) with mass spectrometry (MS) that was employed here combines quantitative analysis with unambiguous identification of captured target molecules in a single workflow [1-3]. SPRi allows to monitor binding affinities and kinetics in real-time and at very low amounts (femtomole range). SPR surfaces are reusable due to regeneration steps between different injections, which allow multiple measurements at different concentrations and, if necessary, various conditions. To identify the molecules that were captured on an SPR surface, matrix-assisted laser desorption/ionization mass spectrometry (MALDI MS) and peptide mass fingerprinting were implemented. By performing SPR and MALDI MS separately, as usually done [4-6], the throughput and sensitivity would be seriously limited due to the need for target elution, which often leads to sample loss and contaminations. Therefore, performing MALDI MS measurements directly on the SPR chip is beneficial [7,8]. We have previously shown that the identity of interacting proteins not only from pure target solutions but also from cellular lysates can be established [9]. The only other similar example reported is the study by Musso *et al*., which demonstrated the detection of α-amylase from human saliva by antibody-arrayed SPRi surfaces [10]. To the best of our knowledge, screening for RNA binding proteins from cell lysates or specific compartments in e.g. mitochondria, with SPRi-MALDI MS has never been performed before.

For unambiguous characterization of captured proteins, mass spectra of high quality are required. In the past, either self-digested peptides of trypsin dominated mass spectra and/or a small number of peptides were obtained from a single, purified injected protein [10-12]. Increasing the efficiency of on-chip proteolysis is therefore essential. To perform both analyses directly on the same surface, working with a MALDI MS compatible SPR chip surface is necessary. Thus, our study also includes improved on-chip digestion, yielding a sufficient number of peptide peaks for confident classification. MS and MS/MS of low protein amounts in the fmol range from arrayed spots were obtained, resulting in increased protein identification reliability.

For this study, we worked with the mitochondrial intron *Sc*.ai5γ from *Saccharomyces cerevisiae* (*Sc*.). It belongs to the class of group IIB introns, natural catalytic RNAs, which share mechanistic and structural similarities with the spliceosome [13-15]. While *in vitro* self-splicing of group II introns can be initiated by high concentrations of divalent metal ions [16,17], *in vivo* the ribozyme functions presumably as a stable ribonucleoprotein (RNP). Structurally, group II introns are highly conserved and display six domains (DI – DVI) radiating from a central wheel [18]. Domain I (DI) is the largest domain, which contains essential binding sites for recognizing its exons and its natural protein cofactor - Mss116.

Mss116 is a DEAD-box helicase and a member of a large ATP-dependent enzyme family involved in various aspects of RNA metabolism [19,20]. Functionally, Mss116 acts not only as an on-pathway intermediate chaperon, preventing kinetic traps during the RNA folding process, but it is also involved in a number of other activities such as strand annealing, conformational switching, and charge neutralization [21-27]. Mss116, as the only natural identified cofactor for *Sc*.ai5γ maturation, is encoded in the nucleus and is transported to the mitochondrial matrix via a signal-peptide, where it can bind to group I and II intron RNAs and trigger their folding and subsequent splicing [22]. The mechanism of Mss116-mediated splicing includes resolving misfolded structures by RNA unwinding and stabilizing correctly folded on-pathway intermediates. Fedorova *et al*. showed that the protein directly stimulates the RNA folding in an ATP-independent manner. Still, ATP is required for the protein turnover - the dissociation from incorrectly formed complexes and rebinding unfolded RNA [25]. However, *in vitro* studies indicate incomplete splicing efficiency, which is in line with our hypothesis that other protein cofactors play a key role in the complete *Sc*.ai5γ maturation [23,28,29].

Although the interaction between *Sc*.ai5γ and Mss116 appears non-specific, the protein function, as an RNA cofactor, has been shown in the past by northern blotting [22]. Equilibrium binding assays by Halls *et al*. [30] demonstrated that Mss116 is able to interact with other classes of introns as well, namely group I and group II introns, with dissociation constants (*K*_D_) in the low nanomolar range, but the association and dissociation kinetic constants (k_on_ and k_off_) were not determined. Mss116 is the only reported protein cofactor involved in *Sc*.ai5γ maturation, pre-mRNA splicing and ribosome assembly. However, the role of other mitochondrial proteins in *Sc*.ai5γ maturation remains unclear. While Mss116 was identified in a screening for nuclear mutants and their ability to grow in the presence or absence of introns [21], we demonstrate the ability of a hyphenated analytical method to discover other proteins that are supposed to be involved in the RNA binding process. With this, we successfully analyzed the lysate of a rough mitochondrial fraction from yeast and identified four previously unknown potential *Sc*.ai5γ-interacting partners.

## 2. Materials and methods

### 2.1. Materials

Trypsin Gold (mass spectrometry grade) was purchased from Promega. DNA linkers, oMGG03 and oMGG05, were ordered at IBA Lifesciences (Göttingen, Germany) and oMGG06 from Microsynth AG (Balgach, Switzerland). All sequences are given in the Supporting Information. A. M. Pyle, Yale University, kindly provided plasmids pJD20 and pHRH108. All chemicals were ordered at the highest available grade from Acros, Fluka, ABCR-Chemicals, or Merck. All solutions for SPR measurements were prepared using DNase, RNase, Protease free water (Molecular Biology Reagent, Sigma-Fine Chemicals).

### 2.2. RNA preparation

The group IIB intron *Sc*.ai5γ DNA, flanked by 41 nucleotides (nt) and 14 nt long 5’- and 3’-exon respectively, was generated by polymerase chain reaction (PCR) amplification using the plasmid pJD20 as a template [31,32], and primers oMGG01 and oSPA37, yielding finally plasmid pMGG01. The pMGG01 DNA construct was *in vitro* transcribed using an in-house produced T7 RNA polymerase. The RNA transcript (referred to as ex^41^-*Sc*.ai5γ-ex^14^ – superscript numbers indicate the number of nucleotides from the flanking exons) was purified on an 8 % denaturing polyacrylamide gel, extracted by electroelution in Tris-borate-EDTA (TBE, pH 8.3) buffer, ethanol precipitated and finally dissolved in distilled water (adapted from Gallo *et al*.) [33].

### 2.3. Protein production

The mitochondrial protein Mss116 (UniProt identifier P15424) DNA was derived from plasmid pHRH108 [34]. The full-length Mss116 DNA, lacking the mitochondrial localization sequence, was delivered in a commercially available pET-SUMO champion vector (Invitrogen), which contains an N-terminal His(6) tag, followed by a SUMO tag sequence. Protein expression and purification were performed as described [23,35], with the following modifications: The protein was expressed in BL21 cells. The bacterial cells were resuspended in a lysis buffer (50 mM Tris-HCl pH 7.5, 500 mM NaCl, 10 mM imidazole, 10 % sucrose, 5 mM β-mercaptoethanol, 1 % IGEPAL-630, and 10 % glycerol). Cell lysis was performed by shock freeze in liquid nitrogen, thawing on ice, and sonication. The lysate was centrifuged at 20’000 *g* at 4 °C for 30 min. The supernatant was filtered and purified by gravity flow Ni-NTA affinity chromatography. The chromatographic columns were filled with 2.5 mL Nickel-Chelating-Resin 50 % slurry in 20 % ethanol (G-Bioscience), washed with 10 column volumes of lysis buffer, and finally eluted in 2 mL fractions in a lysis buffer containing 250 mM imidazole. Mss116 protein was further purified by fast protein liquid chromatography (FPLC) Superdex 200 (GE Healthcare), the protein purity was checked by sodium dodecyl sulfate polyacrylamide gel electrophoresis (SDS PAGE) (detailed results are in the Supporting Information Fig. S1) and MS. Aliquots were stored at −20 °C in a storage buffer (50 mM Tris-HCl pH 7.5, 500 mM NaCl, 1 mM DL-dithiothreitol (DTT) and 10 % glycerol). The protein activity was measured by an ATPase activity kit (Sigma-Aldrich) as described by the manufacturer.

### 2.4. Preparation of rough mitochondrial fractions

The protocol was adapted from Glick and Pon [36]. Harvested yeast cells were washed with water and collected by centrifugation at 4’000 *g* for 5 min at 25 °C. The pellet was resuspended in 100 mM Tris-HCl buffer (pH 8.4) and 10 mM DTT for 10 min at 30 °C and centrifuged at 2’000 *g* for 5 min at 25 °C. After washing with 1.2 M sorbitol and centrifuging for 5 min at 4’000 *g* at 25 °C, the pellet was resuspended in 40 mL buffer containing 1.2 M sorbitol, 20 mM KH_2_PO_4_ (pH 7.4) and 3 mg/g Zymolyase and incubating for 60 min at 30 °C in a water bath. All the subsequent procedures were performed at 0-4 °C. The suspension was centrifuged for 5 min at 2’000 *g*, the supernatant removed, and 100 mL homogenization buffer (10 mM Tris, pH 7.4, 1 mM EDTA, 0.2 % BSA, 1 mM PMSF, and 0.6 M sorbitol) was added to the pellet and mixed thoroughly. The suspension was homogenized using a Dounce homogenizer and then centrifuged for 5 min at 4’000 *g*. The supernatant was saved and additionally centrifuged for 12 min at 7’000 *g*. The resulting mitochondrial pellet was resuspended in 1 mL buffer containing 500 mM KCl, 80 mM MOPS (pH 6.9), 10 mM MgCl_2,_ and 0.5 % Triton-X.

### 2.5. On-plate digestion optimization

To improve the quality of the on-chip tryptic digestion of captured proteins after SPRi measurements, bovine serum albumin (BSA) was used in varying amounts (1 pmol, 100 fmol, and 10 fmol) as a control protein. To focus on the influences of added solutions and chemicals, BSA was spotted on a plate without extra immobilization. Furthermore, all steps were performed without explicitly mixing the solution on the plate. For reduction, DTT and Tris(2-carboxyethyl)phosphine hydrochloride (TCEP) were used at different concentrations (50 mM, 10 mM, 5 mM, 1 mM, 0.5 mM, 0.1 mM and 0.01 mM in 25 mM NH_4_HCO_3_, pH 8.5). 1 µL of the reducing agent was spotted on top of the BSA spots and allowed to react for 1 h, 40 min, 30 min, 20 min, or 10 min at 37 °C. For alkylation, 1 µL of iodoacetamide solution of varying concentrations (5 mM, 1 mM, 0.5 mM, and 0.1 mM) was spotted and incubated for 1 h at 25 °C in the dark. 1 µL of reducing agent was added with the same concentration as the used iodoacetamide (in 25 mM NH_4_HCO_3_ buffer, pH 8.5) to quench the excess of iodoacetamide. Subsequently, 1 µL of trypsin solution (5 ng/µL) was added to each spot and incubated at 37 °C for 10 min. Finally, the spots were dried at 37 °C, and the matrix solution (7 mg/mL α-Cyano-4-hydroxycinnamic acid (α-CHCA) in 50 % acetonitrile/water with 0.3 % trifluoroacetic acid (TFA)) was spotted.

### 2.6. Surface Plasmon Resonance Imaging

All SPRi measurements were carried out using a commercial SPRi-Plex II instrument and standard gold SPRi slides for amine or biotin coupling, respectively (Horiba France, Palaiseau, France). RNA and BSA, as controls, were immobilized by spotting 0.3 µL (0.1 mg/mL) manually on the chip in a 4 x 4 spot array pattern. The blocking buffer consisted of 1 M aqueous ethanolamine (pH 9.0) or 10 µg/mL biotin, respectively. For joining the SPRi slide with the surface of a glass prism with a high refractive index, a thin film of index-matching oil was used to connect both surfaces. SPRi experiments were performed at 25 °C with buffer containing 500 mM KCl, 80 mM 3-(N-morpholino)propanesulfonic acid (MOPS), pH 6.9, 10 mM MgCl_2,_ and 10 µM ATP, respectively, with a flow rate of 50 µL/min and a 150 µL injection volume. For surface regeneration between measurements, 100 mM glycine hydrochloride (H_2_NCH_2_CO_2_H HCl, pH 2.0) was used.

Measured kinetic curves were evaluated using the Scrubber Gen2 software. For calculating the kinetic and binding constants *k*_on_, *k*_off_, and *K*_D_, a classical Langmuir model was applied.

In preparation for the immobilization step, RNA was hybridized with short DNA linkers (oMGG03, oMGG05, or oMGG06) in distilled H_2_O in a 1:1 ratio by heating for 3 min at 90 °C and then slowly cooling down to 25 °C.

### 2.7. MALDI MS

For MALDI MS measurements, a tryptic digestion was performed directly on the chip surface after being removed from the SPRi instrument and air-dried. The disulfide bonds of the target proteins were reduced for 1 h at 37 °C with 500 pmol of aqueous TCEP in 25 mM ammonium bicarbonate (NH_4_HCO_3_), pH 8.5. The addition of 25 mM NH_4_HCO_3_ solution prevents the sample from drying out. Trypsin (5 ng/µL in 10 mM NH_4_HCO_3_, pH 8.3) was spotted (0.3 µL) onto the region of interest and kept in a humid atmosphere at 37 °C for 10 min. Finally, after the spots had dried and digestion had stopped, a MALDI matrix was applied (0.1 µL of 7 mg/mL α-CHCA in 50 % acetonitrile/water with 0.3 % TFA). For MALDI MS measurements, the SPRi slide was mounted directly into a custom-made SPRi-MALDI adapter. Mass spectra were obtained with an Ultraflex II MALDI-TOF (Bruker Daltonics, Bremen, Germany), which is equipped with a “smartbeam” laser (337 nm UV laser based on a modified solid-state laser) working with a 66.7 Hz repetition rate, using the following parameter settings: positive ion reflectron mode, acceleration voltage: 25 kV, delayed extraction time: 20 ns, laser fluence: just above the desorption/ionization threshold, number of laser shots: 1’500-2’500 (averaged and acquired at random sample positions). All mass spectra were smoothed, baseline-subtracted, and externally calibrated with a standard peptide calibration mix I (LaserBio Labs), which was also spotted onto the SPRi slide. Tryptic peptides were identified with the Biotools software (Bruker Daltonics) and Mascot software (Matrix Science Limited).

### 3. Results and discussion

### 3.1. Optimization of on-chip digestion

In order to unambiguously identify proteins, on-chip proteolysis first needed to be improved. As previously described [9], we optimized the concentration of trypsin (5 ng/µL) as well as the incubation time (10 min), salt concentration (10 mM NH_4_HCO_3_, pH 8.3), and temperature (37 °C). Because of the relatively small number of tryptic peptides generated (e.g. 8 peptides from a purified protein), which leads to a low sequence coverage (e.g. 12.3 %) [9], we have further improved the on-chip proteolysis with BSA as a model protein. First, the influence of reduction prior to the addition of trypsin was studied. To the best of our knowledge, mainly two reagents are reported as reducing agents in tryptic digests, DTT and TCEP [37-40]. While keeping the reaction time constant, the concentration of the reducing agents was varied between 0.01 to 50 mM. Optimal results were obtained at 0.5 mM for both reducing agents (Supplementary Table S2). Higher concentrations of reducing agents decreased the sensitivity, as many of the peptides detected at lower concentrations of reducing agent could no longer be observed. This observation indicates that high reducing agent concentrations have a strong suppressing effect. In addition, concentrations below 0.5 mM reduced the quality of the digest, with a substantial decrease in sequence coverage (e.g. from 62.1 % to 27.5 % for 100 fmol BSA with 0.5 mM and 0.1 mM TCEP, respectively) (Supplementary Table S2). The comparison between the peptide mass fingerprint analysis using DTT or TCEP, respectively, clearly shows that TCEP reacts significantly better (e.g. 68.0 % sequence coverage for 100 fmol BSA) than DTT (e.g. 24.7 % sequence coverage for 100 fmol BSA) by a factor of more than two in sequence coverage (Fig. 1 A). Furthermore, compared to tryptic digestion without reduction, the reduction with TCEP doubles or even triples the sequence coverage, depending on the amount of protein (Fig. 1 A). It is noteworthy that we achieved a proteolytic digest of only 10 fmol BSA yielding 42.7 % sequence coverage. Consequently, 0.5 mM TCEP was chosen for all further experiments.

**Fig 1.**
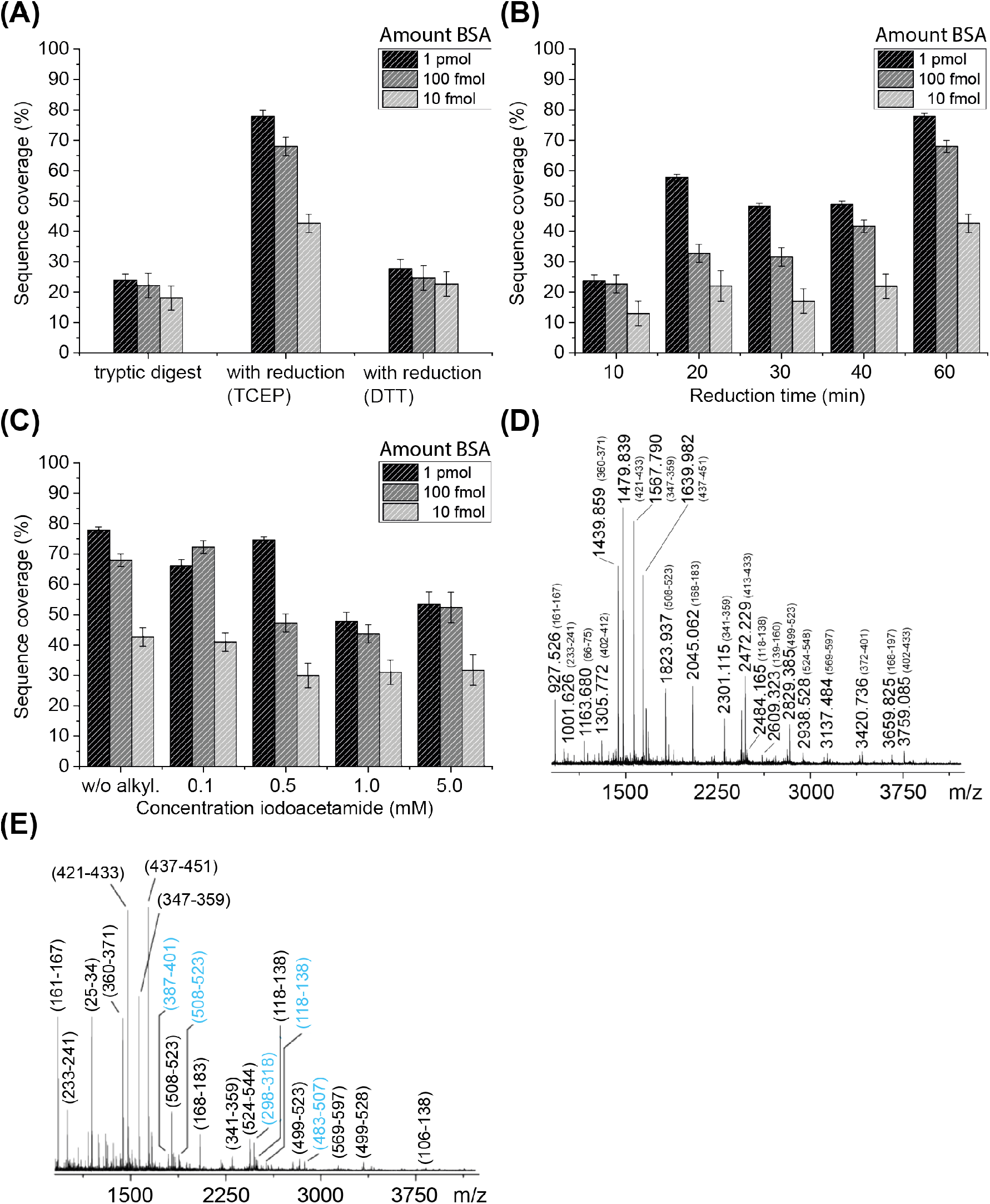
Comparison of sequence coverage from on-plate digested BSA (1 pmol, 100 fmol and 10 fmol) with varying conditions. **(A)** Tryptic digestion with and without reducing agent (0.5 mM TCEP and DTT, respectively, for 1 h). **(B)** Percentage of sequence coverage from tryptic digestion with reduced BSA (0.5 mM TCEP) at different time points. **(C)** Effect of the alkylation reagent iodoacetamide on the sequence coverage at different concentrations (w/o alkyl. = without alkylation) after reduction (0.5 mM TCEP, 1 h). **(D)** Mass spectra of reduced (0.5 mM TCEP, 1 h) and digested BSA (100 fmol) in the absence and the presence of alkylation agent (small inlet, 0.1 mM iodoacetamide). **(E)** Mass spectra of reduced (0.5 mM TCEP, 1 h) and digested BSA (100 fmol) in the presence of alkylation agent (0.1 mM iodoacetamide). Fragments are labeled with their sequence range in parentheses. Alkylated peptides are highlighted in blue. For clarity, not all identified peptides are labeled.

Second, the influence of reduction over time was investigated. Minimizing the duration for reduction from 60 to 10 min induced a steady decrease in sequence coverage (Fig. 1 B). A reduction to 10 min resulted in a similar low sequence coverage as for the tryptic digestion without the reduction (between 10 to 25 %).

It is known that reduced disulfide bonds are often alkylated to prevent the reformation of disulfide bonds and thus to refold [41]. Therefore, we investigated whether an additional on-plate alkylation step further improves the coverage of the protein sequence. For this purpose, iodoacetamide was applied to increase the concentration from 0.1 to 5 mM, while keeping reduction conditions, like temperature and duration time, constant (Fig. 1 C). The use of this alkylation reagent has a similar impact on the digest as the reducing agents. The higher the concentration of iodoacetamide, the lower the sequence coverage, especially at lower BSA amounts. Low-intensity peaks are suppressed with higher concentrations of alkylation reagent. Overall, no new fragments were observed (Fig. 1 D and 1 E). Since no significant increase in sequence coverage was achieved by the addition of an alkylation step, but the experimental time almost doubled, alkylation was omitted in further experiments. These results show that we are able to overcome the difficulties of identifying proteins that occur in very small amounts, i.e. 100 – 10 fmol, by peptide mass fingerprint analysis.

### 3.2. Binding of ex^41^-Sc.ai5γ-ex^14^ to Mss116

To date, only the affinity of Mss116 to *Sc*.ai5γ has been determined [30], but the binding and unbinding kinetics remain unknown. Initially, SPRi measurements were performed to characterize the association and dissociation rate constants (*k*_on_ and *k*_off_) of Mss116 to the intact ex^41^-*Sc*.ai5γ-ex^14^. These constants give insight into how fast and specific the interaction between both molecules occurs and thus provide a better understanding of the biological system and exact mode of action. In a second step, we used SPRi-MALDI MS to identify additional RNA-binding proteins from a lysate of a rough mitochondrial fraction, which required improved on-chip proteolysis.

To attach the RNA onto a surface, two different immobilization strategies were applied: a biotinylated DNA oligonucleotide was hybridized either to domain IV (DIV) or to the 5’exon of ex^41^-*Sc*.ai5γ-ex^14^ (Fig. 2 A). Hence, we examined whether the binding properties of Mss116 to the RNA are influenced due to a structurally different conformation and accordingly identify the better preferable immobilization position for an optimal interaction. Injections of Mss116 at various concentrations (5-135 nM) showed a slow dissociation, which does not reach the baseline level during the measuring time of 650 sec (Fig. 2 B and Table 1). In agreement with previous investigations [34,42], the measured binding rates indicate a rapid binding of Mss116 (with a fast association rate constant) and a prolonged release (Fig. 2 B and Table 1).

**Fig 2.**
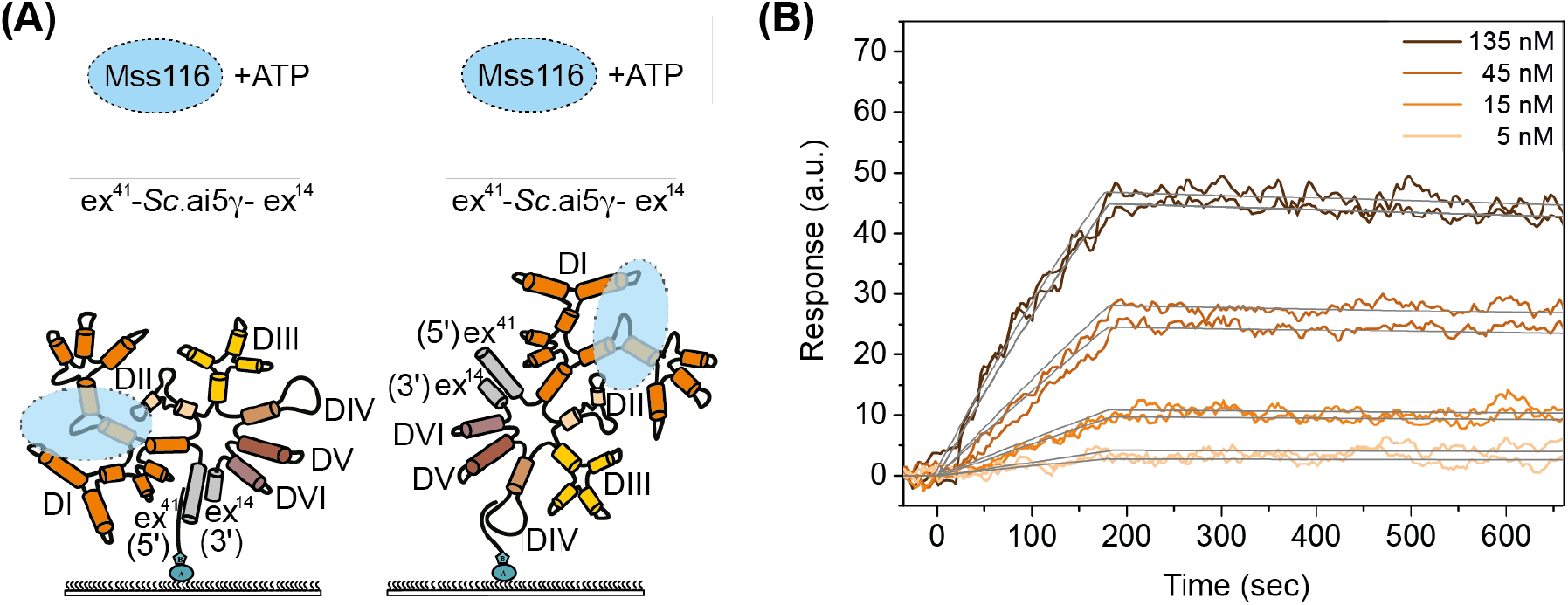
**(A)** Scheme of the secondary structure of ex^41^-*Sc*.ai5γ-ex^14^ indicating the proposed binding site for Mss116 (blue) at a central three-way junction (κ-ζ) within DI (orange). The different immobilization approaches via the 5’exon (left) and DIV (right) are shown. **(B)** SPRi measurement of the group IIB intron ex^41^-*Sc*.ai5γ-ex^14^ in the presence of ATP (10 µM) in running buffer and a concentration series of Mss116 (5-135 nM). Calculated *K*_D_: 8.4 ± 0.7 nM (fits in grey).

**Table 1.** Association (*k*_on_) and dissociation (*k*_off_) rate constants and *K*_D_ of the interaction between ex^41^-*Sc*.ai5γ-ex^14^ and its cofactor Mss116 in the presence or the absence of ATP for both immobilization schemes (DIV and 5’exon). Standard deviations derive from two to four independent technical replicates.

| **Immobilization tag and position** | **ATP** | **$k_{\text{on}}$ ($\text{M}^{-1}\text{s}^{-1}$)** | **$k_{\text{off}}$ ($\text{s}^{-1}$)** | **$K_D$ (nM)** |
| --- | --- | --- | --- | --- |
| **biotin – DIV** | No | $(5.76 \pm 0.07) \times 10^4$ | $(3.11 \pm 0.04) \times 10^{-4}$ | $5.40 \pm 0.06$ |
| **biotin –DIV** | Yes | $(1.20 \pm 0.10) \times 10^4$ | $(9.90 \pm 0.50) \times 10^{-5}$ | $8.40 \pm 0.70$ |
| **biotin – 5'exon** | Yes | $(3.00 \pm 0.10) \times 10^4$ | $(2.70 \pm 0.10) \times 10^{-4}$ | $9.10 \pm 0.30$ |
| **amine –DIV** | Yes | $(6.51 \pm 0.05) \times 10^4$ | $(1.35 \pm 0.01) \times 10^{-4}$ | $2.07 \pm 0.02$ |

Omitting ATP (10 µM) from the running buffer was found to have only little effects on the binding strength, with a *K*_D_ value in the low nanomolar range (Table 1 and Fig. S2) as well as unaffected on- and off-rates. Variations in *K*_D_ by a factor of 3 to 4 obtained from biophysical experiments are within the tolerance range. Our findings on the low impact of ATP are in good agreement with previous results, where it was shown that the presence of ATP has no influence on the protein-RNA binding strength [23]. In contrast, it was reported that the release of Mss116 from the RNA is ATP-dependent [25,43]. Therefore, the following experiments were performed in the presence of ATP. Nevertheless, we could not observe the complete dissociation of the protein from ex^41^-*Sc*.ai5γ-ex^14^, and under our experimental conditions, the protein remains bound to the target molecule.

We tested two different immobilization approaches (DIV and 5’exon immobilization) to ensure that Mss116 binding is not hindered by the surface-immobilization, despite the binding site is believed to be in DI. The determined *K*_D_-values for both approaches are very similar and within the experimental error (Table 1, Fig. S2, and S3). No significant changes in the binding of Mss116 were observed when the biotin tag was replaced with an amine tag. The tag exchange was necessary for the DNA linker hybridizing to DIV and the target immobilization on a protein-free SPRi slide (Table 1 and Fig. S4). The results demonstrated that neither DIV nor the 5’exon is involved in Mss116 binding, and DI is not concealed in both of the immobilization approaches, which is in line with previous findings that DI is the preferred counterpart for Mss116 [29]. Additionally, the measured binding rates indicate a rapid binding of Mss116 (with a fast association rate constant) and a prolonged release.

### 3.3. SPRi-MALDI MS analysis of rough mitochondrial lysate from baker’s yeast

Having shown that immobilization by either DIV or via the 5’exon does not affect Mss116 binding to ex^41^-*Sc*.ai5γ-ex^14^, we proceeded with SPRi-MALDI MS measurements and used the surface-immobilization via DIV. Although it is known that members of the DEAD-box helicases are often involved in multiple processes, such as mitochondrial intron splicing and transcription elongation [44], the interaction between *Sc*.ai5γ and other potential interactors has been poorly investigated. The hypothesis that group II introns and the spliceosome are evolutionarily related is well established. Following the theory of Galej and coauthors [45] that spliceosomal proteins gradually replaced intronic domains during the evolution from bacteria to eukaryotes, the presence of more than one cofactor for the mitochondrial group II intron is very likely. We assume that besides Mss116, other regulating factors participate in the splicing process of *Sc*.ai5γ. To support this hypothesis, we combined SPRi with MALDI mass spectrometry to screen and identify new potential intron binders. This method allows also the detection of interacting analytes in real time and the determination of association and dissociation constants.

Firstly, on-chip tryptic proteolysis needed to be optimized (section 3.1). Secondly, by performing SPRi-MALDI MS analysis, we faced another challenge. Although immobilization via biotin or amine tagged RNA resulted in only marginal differences in binding constants, it significantly affected the MALDI MS analysis. Biotinylated RNA is immobilized on SPRi slides coated with *ExtrAvidin*, a modification of avidin with high affinity and specificity to biotin. However, this caused too many disrupting, intense peaks in the mass spectrum (Fig. S5 A and B) coming from the *ExtrAvidin* protein coating when spots of interest are digested with trypsin. The replacement of the enzyme trypsin with AspN, to generate *ExtrAvidin*-peptides with higher m/z values did not improve the situation. In this case, autolysis products of AspN dominated the entire mass spectrum (Fig. S5 C and D), even when the same low concentrations as for trypsin were used. Finally, to avoid interfering peaks, we used SPRi slides in the absence of any protein functionalization. We chose N-hydroxy succinimide (NHS) ester functionalized surfaces, which require free amine groups on the molecule for immobilization. Obtaining high-quality mass spectra from on-chip proteolysis of captured analytes (in the low fmolar range) from arrayed SPRi slides is considered challenging. By using our optimized workflow, after the immobilization of the amine-tagged ex^41^- *Sc*.ai5γ-ex^14^ (DIV), captured Mss116 could be identified after the injection of the pure protein, with a reasonably high number of fragments and a sequence coverage of 68 % from only 9.4 fmol (Fig. 3 bottom and Fig. S6).

**Fig 3.**
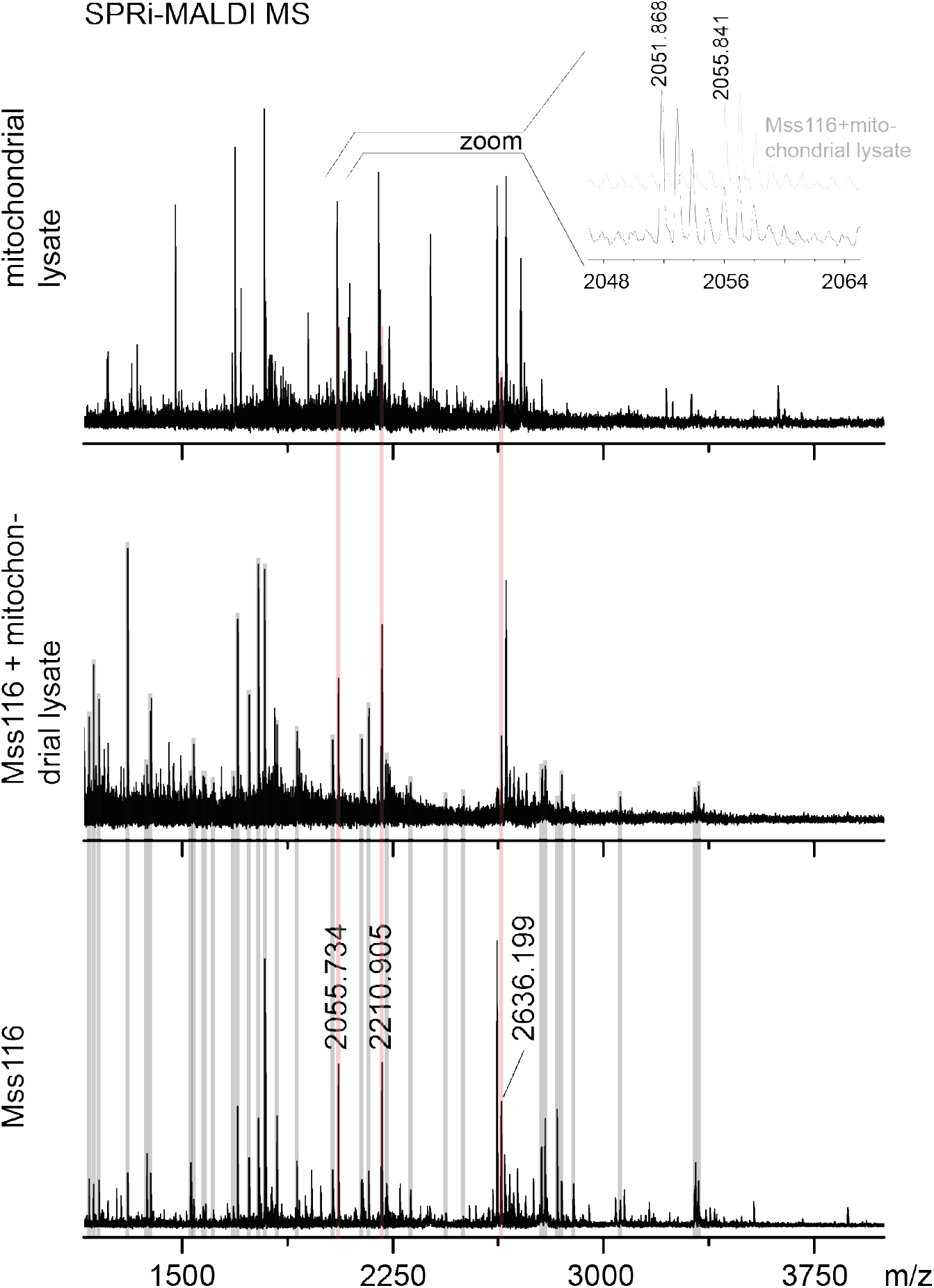
On-chip MALDI mass spectra after SPRi measurements and on-chip proteolysis (with reduction and trypsin as proteolysis enzyme) after **(top)** the direct injection of a rough mitochondrial fraction lysate, **(middle)** the injection of 250 nM Mss116 (causing saturation in the SPR sensorgram) and mitochondrial fraction without regeneration in between, and **(bottom)** the injection of Mss116 (135 nM). Grey bars mark the same peptides found in the spectra **middle** and **bottom**, while light red highlighted peptides were found in all three experiments. A complete assignment of the peptide mass fingerprint from Mss116 **(bottom)** can be found in Fig. S6 (Supporting Information). Assignments of the peaks from the lysate **(top)** are given in Table 2.

To investigate if proteins other than Mss116 interact with ex^41^-*Sc*.ai5γ-ex^14^ we applied a rough mitochondrial lysate from yeast to the immobilized RNA. The MALDI MS experiment was performed after a single injection of the lysate and based on real-time detected plasmon curves, i.e. if an interaction is detected/not detected. The mass spectrum of the on-chip tryptically digested spots of interest revealed a high number of peptides (Fig. 3 top). A comparison between the peptide mass fingerprint spectrum of Mss116 (Fig. 3 bottom) and the mitochondrial fraction (Fig. 3 top) shows distinct differences, i.e. many peptides with different m/z values were observed, indicating that proteins other than Mss116 bind to ex^41^-*Sc*.ai5γ-ex^14^. Only three relatively low intensity peaks were observed, e.g. the signal at m/z 2055.841 (Fig. 3 top, zoomed part), indicating binding of Mss116. A Mascot database query of the fragments shows that several other proteins interact with the target RNA as well. Based on the peaks with a signal-to-noise ratio higher than 1.5 and a mass deviation of less than 190 ppm (as achieved with on-chip digested Mss116), we identified four distinct proteins, namely the pre-mRNA-splicing factor 8 (Prp8), DEAD-box protein 8 (Dbp8), mitochondrial ribosomal protein 13 (Mrp13) and a protein known as Cullin-3. The identity of the last two proteins was further verified by MS/MS measurements of selected ions. The MS/MS spectrum of ion m/z 1791.704 (Fig. 4) confirmed the amino acid sequence AKLDEFLIYHKTDAK originating from Mrp13. The fragmentation of the ion m/z 2198.940 supports the presence of Cullin-3 (Fig. S7).

**Table 2.**
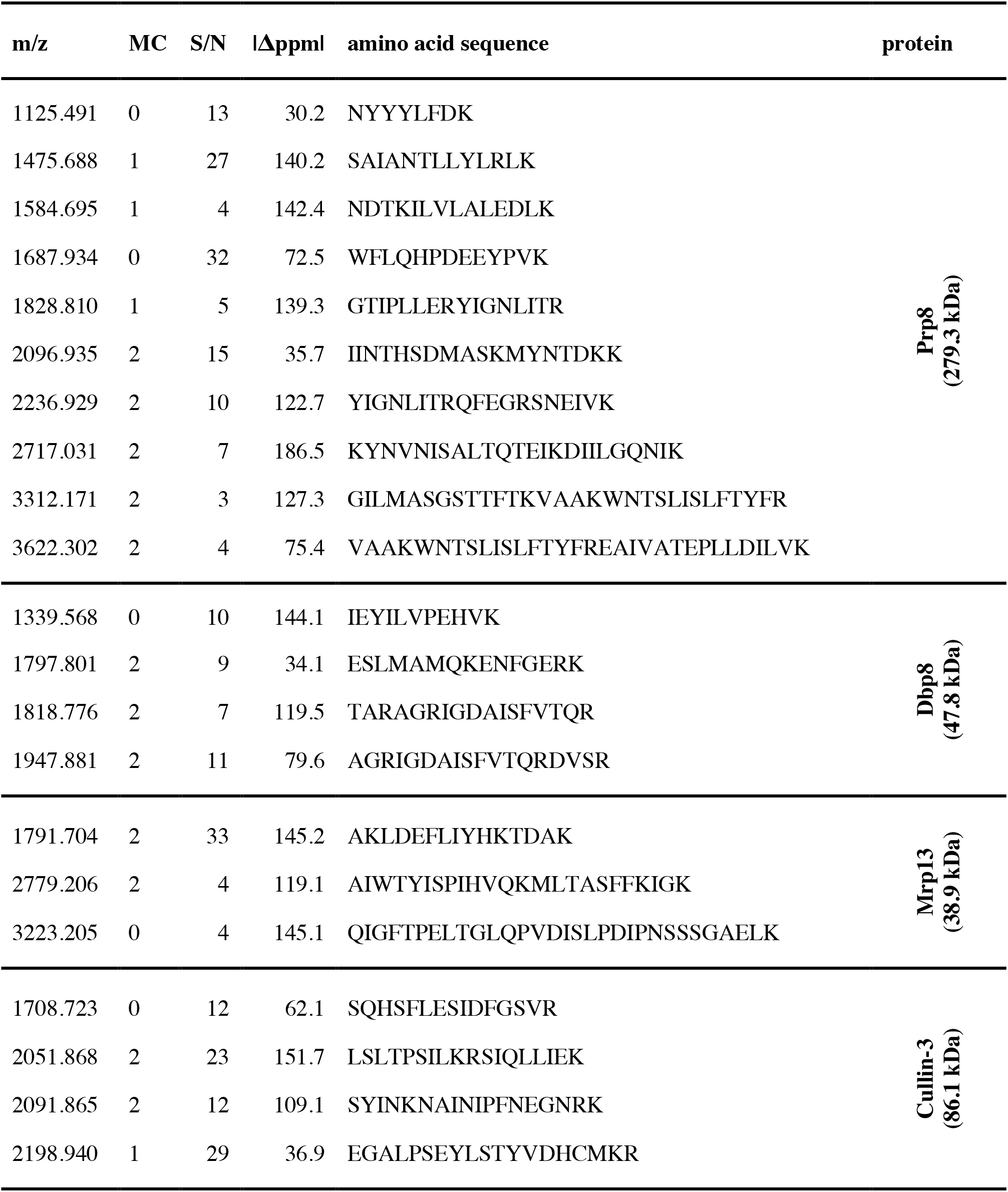
Captured and identified peptides after the injection of a rough mitochondrial fraction lysate and on-chip proteolysis using MALDI MS (compare Fig. 3 top). Peptides with their m/z value, number of missed cleavages (MC), signal-to-noise ratio (S/N), deviation from theoretical m/z (Δppm), and amino acid sequences are grouped into each identified protein resulting from a Mascot database request and peptide mass fingerprint.

**Fig 4.**
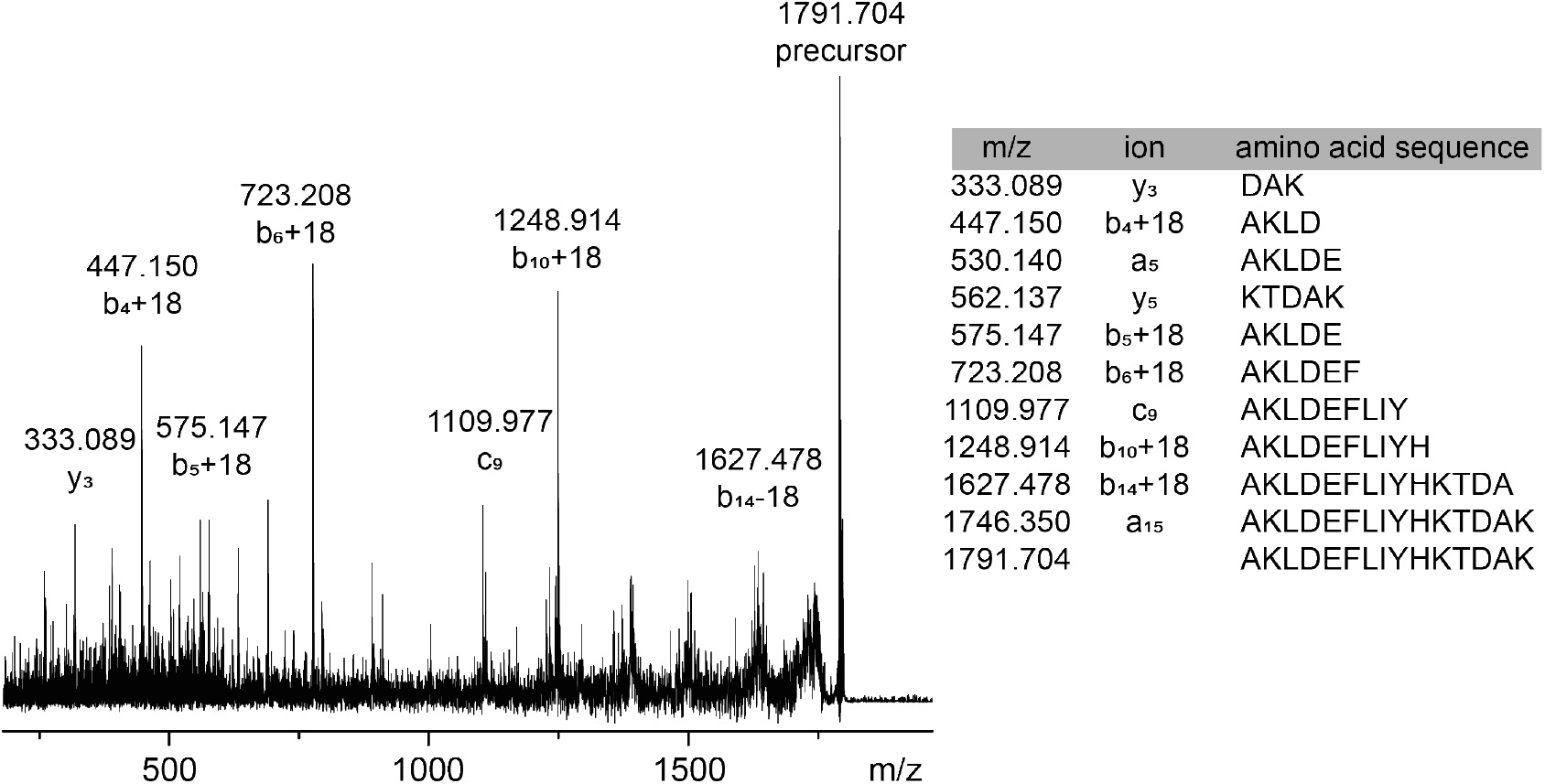
SPRi-MALDI MS: On-chip MS/MS measurement of the peptide m/z 1791.704 (from Mrp13), which was detected after injecting a rough mitochondrial fraction lysate. A complete list with identified fragments (m/z values, cleavage site of the peptide (“ion”) and positions of involved amino acids, and the amino acid sequence) is given in the table to the right.

To exclude non-specific adsorption resulting from the SPRi slide surface and thus false-positive signals, we used reference spots (without immobilized ex^41^-*Sc*.ai5γ-ex^14^) as controls. Here, no increasing signal in plasmon curves after adding the lysate or peptides from MALDI MS measurements was observed (data not shown).

By screening a rough yeast mitochondrial lysate with SPRi-MALDI MS, we identified four new potential binders of the group II intron *Sc*.ai5γ, in addition to the well-known splicing cofactor Mss116. In the following section, we explain why these four proteins, namely Dpb8, Prp8, Mrp13, and Cullin-3, can bind to *Sc*.ai5γ. The ribosomal protein Dbp8 shares sequence similarities with Mss116, e.g., both are members of the DEAD-box family, named after the highly conserved amino-acid sequence D-E-A-D (asp-glu-ala-asp). This so-called Walker motif is required for ATP binding and hydrolysis. In addition, Dbp8 is an RNA helicase known to participate in synthesizing the small ribosomal subunit (18S rRNA) and interacting with the target RNA via five conserved motifs (Ia, Ib, II, V, and VI). [46,47]. In contrast to Dbp8, the pre-mRNA-splicing factor 8 (Prp8) consists of domains analogous to the intron-encoded proteins (IEP), which assist in splicing [48]. Introns that lack such an IEP are supported by ATP-dependent helicases, like Mss116. Prp8 is part of the yeast spliceosome and the key regulator of intronic RNA splicing in the nucleus. The large domain of Prp8 interacts with both splice sites and the essential branch point by forming the catalytic core of the RNAP complex, very similar to the maturase domain in IEPs [45,49-51] Mrp13 is part of the small subunit of the mitochondrial ribosome, hence its name [52-54]. Cullin-3 is described as a scaffold protein involved in the degradation-related ubiquitylation pathway [55-57]. Similar to Mss116, both proteins carry hydrophilic residues that facilitate RNA binding. All four newly identified proteins comprise RNA-binding properties, either identical to Mss116 or to an IEP, that enable them to recognize RNA motifs present in *Sc*.ai5γ. Among these binders, Mrp13 is the only mitochondrial protein. The other three proteins are localized in the nucleus. The way these nuclear proteins can enter the mitochondria and whether they are localized remains unclear. The three nuclear proteins do not carry any signal peptide for the directed transport into the mitochondria. However, there are possible mechanisms known as ADP/ATP carrier via TIM22 complex, stress, or passive internalization to deliver cellular or nuclear proteins into mitochondria rather than the signaling peptide pathway [58,59]. It has been already shown that non-mitochondrial proteins and proteins derived from species different than yeast trigger the splicing in *Sc*.ai5γ *in vitro* [23]. Solem *et al*. demonstrated that the cytoplasmic ATP-dependent helicase DED1 and Cyt-19 in *Neurospora crassa*, both members of the DEAD-box family, stimulate *Sc*.ai5γ splicing *in vitro*. Suggestively, this is due to sequence and functional similarities to Mss116, and thus, these proteins bind similar RNA motifs. The here found RNA-binding proteins are known to be multifunctional, and therefore it is not surprising that they can bind different kinds of RNAs [30,60,61]. Similar to Mss116, which interacts with group I and II introns, the identified proteins bind other RNAs and participate in one way or another in the RNA maturation process, as described in detail above. On the other side, group II introns are characterized by a complex secondary and tertiary structure highly conserved among all species. *Sc*.ai5γ comprises structural similarities with other intronic RNAs as well, and its cofactor Mss116 shares universal features with similar functioning RNA-binding proteins [62]. It is possible that the proteins found bind the model-RNA based only on the sequence and structural similarities, with respect to the RNA but also regarding the cofactors Mss116 and IEP. Hence, the identification of four additional *Sc*.ai5γ binding candidates is a first experimental indication of other potential cofactors that may be involved in the group II intron splicing process *in vivo*. Their presence in mitochondria and their biological relevance and interaction with *Sc*.ai5γ *in vitro* and *in vivo* needs further investigation.

For further investigation, we applied epitope blocking to identify potential binding sites between the identified proteins and ex^41^-*Sc*.ai5γ-ex^14^,. Thereby, Mss116 was first injected to saturate its binding site on the RNA that was anchored on the SPR surface. Without regenerating the surface, the yeast mitochondrial lysate was subsequently added. The sensorgram signal increased only marginally compared to the direct injection of the mitochondrial fraction. The mass spectrum (Fig. 3 middle) shows mainly peaks corresponding to Mss116 and only a few others that are different from those of mitochondrial lysate only (Fig. 3 top). There are two possible explanations for this finding: (i) the newly identified binders share the same RNA binding site with Mss116, which cannot be removed because being present in large excess and the binding is not competitive, or (ii) binding of Mss116 to the RNA introduces structural changes that prevent other proteins from binding to their target site, different from the Mss116 binding site. The latter case would imply that the other proteins possibly bind to the group II intron not in the pre-spliced state but rather at a later stage of the splicing reaction. To identify the exact binding regions, further experiments are required. By using different immobilization schemes of ex^41^-*Sc*.ai5γ-ex^14^, we could demonstrate that neither DIV nor the 5’exon is involved in Mss116 binding and that DI is accessible in each of the used immobilization schemes. These results are consistent with the previously proposed Mss116 binding site in *Sc*.ai5γ, namely DI [29]. The obtained results demonstrate that the improved on-chip digestion workflow using SPRi-MALDI MS is a powerful tool for identifying new potential binders.

## 4. Conclusions

RNA-binding proteins are usually grouped into either specific or non-specific ones, where the latter is still poorly understood. How these proteins differentiate between binding sites is still unknown. A possible approach might be to investigate whether the binding is kinetically controlled. This would show differences in association and/or dissociation rate constants, while equilibrium dissociation constants (*K*_D_) may have similar values. The demand for screening and high-throughput analysis tools, including non-targeted approaches, is steadily increasing. The present RNA – protein interaction study demonstrates the potential of SPRi-MALDI MS as a general tool for identifying such interactions by using one of the largest known ribozymes (ex^41^- *Sc*.ai5γ-ex^14^) as a target. Non-targeted, non-covalent interactions are thereby also detected from a cell lysate, a complex mixture of proteins. For determining the identity of binding partners, a complementary technique is adapted. Combining SPRi with MALDI MS in a unified workflow is a powerful tool for identifying target binders from specific cell compartments. So far, similar investigations have only been carried out on known protein-protein interactions [9,10]. The on-chip workflow developed here generates high-quality mass spectra (including better S/N values) and leads to enhanced confidence in protein identification. With the implementation of a reduction step prior to digestion and a proper amount of trypsin (see Materials and methods), we were able to overcome the difficulties of identifying marginal amounts of protein. The results show the first hits of a screening that has been conducted. While further studies will be needed to validate the function and binding strength of the identified proteins, we developed for the first time a screening method for potential RNA binders from a crude yeast extract using SPRi-MALDI MS.

## Authorship contribution statement

The manuscript was written through the contributions of all authors. All authors have approved the final version of the manuscript. UA and MGG contributed equally to this work. UA, MGG, and SZP designed the experiments. MGG and SZP developed different immobilization approaches. MGG produced and purified the functional RNA and its cofactor Mss116. UA performed the SPRi-MALDI MS experiments and the corresponding data analysis.

## Acknowledgment

We thank Dr. Carmelina Petrungaro (group of Prof. Benoît Kornmann, Institute of Biochemistry, ETH Zurich) for kindly cultivating *Saccharomyces cerevisiae* cells and for the introduction into the preparation of mitochondrial lysate. We also thank the Molecular and Biomolecular Analysis Service (MoBiAS, ETH Zurich) for access to the MALDI MS instrument and productive discussions.

## Funding Sources

We thank Horiba Jobin Yvon for the loan of an SPRi instrument and Bruker Daltonics, Horiba Jobin Yvon for jointly funding 1.5 years of a Ph.D. studentship for UA, and Forschungskredit FK 16-083 to MGG. This work was financially supported by UZH (to RKOS) and by the Swiss National Science Foundation, grants # 200020_159929 (to RZ) and 200020_165868/1 (to RKOS).

## Notes

The authors declare no competing financial interest. The original data used in this publication are made available in a curated data archive at ETH Zurich (https://www.researchcollection.ethz.ch) under the DOI 10.3929/ethz-b-000350562.

## Appendix A. Supplementary data

Supplementary data to this article are available and include the amino acid sequence of Mss116; sequences of the DNA oligonucleotides used for different immobilization strategies; the RNA sequence of the full-length group IIB intron with the shortened flanking exons ex^41^-*Sc*.ai5γ-ex^14^, and the primers used for the DNA construct amplification by PCR; SPR measurements in the presence and absence of ATP from different immobilization strategies; MALDI mass spectra showing the effect of *ExtrAvidin*-coated SPRi surfaces; MALDI mass spectrum after SPRi measurement from captured Mss116 with corresponding fragments, MS/MS measurement of the peptide m/z 2198.940 detected after injecting a rough mitochondrial fraction lysate.

## Abbreviations

α-CHCA: α-cyano-4-hydroxycinnamic acid
ATP: adenosine 5’-triphosphate
BSA: bovine serum albumin
dH_2_O: distilled water
DNA: desoxyribonucleic acid
EDTA: ethylenediaminetetraacetic acid
FPLC: fast protein liquid chromatography
His: histidine
IGEPAL-630: detergent (registered trademark)
nt: nucleotide
MALDI MS: matrix-assisted laser desorption/ionization mass spectrometry
MOPS: 3-(N-morpholino)propanesulfonic acid
Ni-NTA: nickel-nitrilotriacetic acid
PCR: polymerase chain reaction
PMSF: phenylmethylsulfonyl fluoride
RNA: ribonucleic acid
Sc: Saccharomyces cerevisiae
SPRi: surface plasmon resonance imaging
SUMO: Small Ubiquitin-like Modifier
TBE: Tris-borate-EDTA
TCEP: tris(2-carboxyethyl)phosphine
TFA: trifluoroacetic acid
Tris: tris(hydroxymethyl)aminomethane

## Notes

https://www.researchcollection.ethz.ch

### Competing Interest Statement

The authors have declared no competing interest.

### Summary of Updates

submission to a different journal, format adapted

