## Supplemental Info for "Screening for Potential Interaction Partners with Surface Plasmon Resonance Imaging Coupled to MALDI Mass Spectrometry"

```
        10         20         30         40         50         60
MGSSHHHHHH GSGLVPRSAS MSDSEVNQEA KPEVKPEVKP ETHINLKVSD GSSEIFFKIK

        70         80         90        100        110        120
KTTPLRRLME AFAKRQGKEM DSLRFLYDGI RIQADQTPED LDMEDNDNIE AHREQIGGSL

       130        140        150        160        170        180
YNDGNRDQRN FGRNQRNNNS NRYRNSRFNS RPRTRSREDD DEVHFDKTTF SKLIHVPKED

       190        200        210        220        230        240
NSKEVTLDSL LEEGVLDKEI HKAITRMEFP GLTPVQQKTI KPILSSEDHD VIARAKTGTG

       250        260        270        280        290        300
KTFAFLIPIF QHLINTKFDS QYMVKAVIVA PTRDLALQIE AEVKKIHDMN YGLKKYACVS

       310        320        330        340        350        360
LVGGTDFRAA MNKMNKLRPN IVIATPGRLI DVLEKYSNKF FRFVDYKVLD EADRLLEIGF

       370        380        390        400        410        420
RDDLETISGI LNEKNSKSAD NIKTLLFSAT LDDKVQKLAN NIMNKKECLF LDTVDKNEPE

       430        440        450        460        470        480
AHERIDQSVV ISEKFANSIF AAVEHIKKQI KERDSNYKAI IFAPTVKFTS FLCSILKNEF

       490        500        510        520        530        540
KKDLPILEFH GKITQNKRTS LVKRFKKDES GILVCTDVGA RGMDFPNVHE VLQIGVPSEL

       550        560        570        580        590        600
ANYIHRIGRT ARSGKEGSSV LFICKDELPF VRELEDAKNI VIAKQEKYEP SEEIKSEVLE

       610        620        630        640        650        660
AVTEEPEDIS DIVISLISSY RSCIKEYRFS ERRILPEIAS TYGVLLNDPQ LKIPVSRRFL

       670        680        690        700        710        720
DKLGLSRSPI GKAMFEIRDY SSRDGNNKSY DYDDDSEISF RGNKNYNNRS QNRDYDDEPF

       730        740
RRSNNNRRSF SRSNDKNNYS SRNSNIY
```

His-tag Linker SUMO-tag Linker mature Mss116

**Scheme S1.** Amino acid sequence of the Mss116 construct as used in this study. The individual parts are indicated by color-coding.

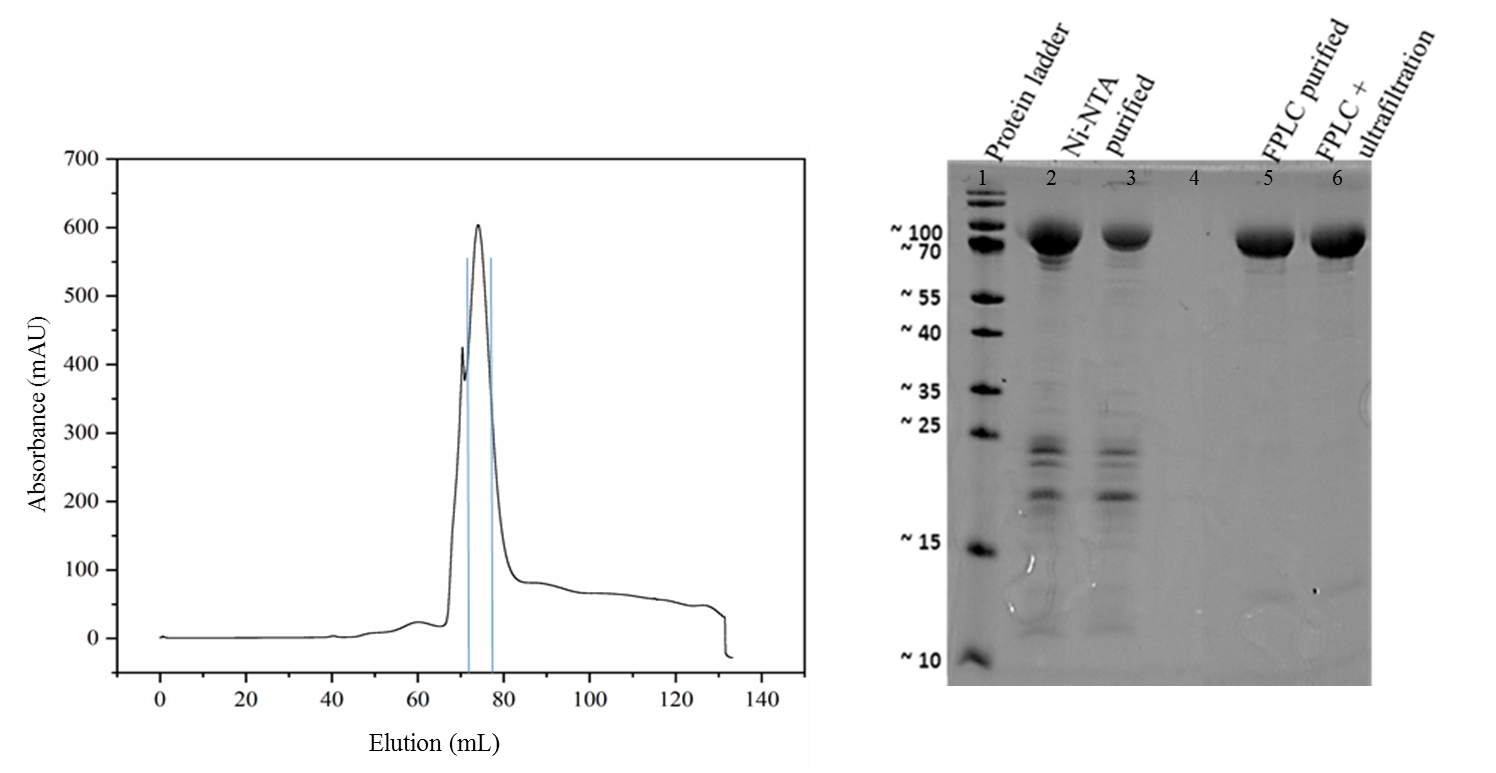

**Fig. S1.** Elution profile of Mss116 purification using FPLC (HiLoad 16/600 Superdex 200 pg,). The main elution peak includes the fractions which were collected (framed in blue) shown on the left and analyzed subsequently on a SDS PAGE. The SDS-PAGE was stained with Coomassie brilliant blue. Indicated are the protein ladder in lane 1, the Ni-NTA purified protein Mss116 in lane 2 and 3 (loaded 5 μL of 520 μM after Ni-NTA purification and 5 μL of the protein sample after dialysis and a concentrating step respectively). Lanes 5 and 6 represent the purified protein Mss116 (15 μl of 73 μM Mss116 was loaded) after FPLC and FPLC, followed by ultrafiltration. After analysis, they were pooled and stored at -20 °C.
**Supplementary Table S1.** DNA oligonucleotides used for the immobilization and the plasmid amplification. oMGG03 and oMGG06 were used to immobilize the group II intron ex^41^-*Sc*.ai5γ-ex^14^ via DIV, oMGG05 for the surface-attachement via the 5'exon. The DNA template for RNA transcription was amplified by PCR with oMGG01 and oSPA37.

| Name of oligonucleotide | Sequence in 5' – 3' direction |
| --- | --- |
| oMGG03 | Cy5-TATATATAAACCTCCTATCGTCCGTACGT-biotin |
| oMGG05 | Cy3-GTAGTAAGTCTCCCCAATAA-biotin 3' |
| oMGG06 | TATATATAAACCTCCTATCGTCCGTACGT(3'C6 Amine) |
| oMGG01 | TCGGAATTCGGGGAGACTTACTACGTGGTGGG- |
| oSPA37 | CGCGCGAAGCTTGATAATACATAGTATCCCG |

5'­GGGCGAAUUGGGGGAGACUUACUACGUGGUGGGACAUUUUCGAGCGGUCUGAAAGUUAUCAUAAAUAAUAUUUACCAUAUAAUAAUGGAUAAAUUAUAUUUUUAUCAAUAUAAGUCUAAUUACAAGUGUAUUAAAAUGGUAACAUAAAUAUGCUAAGCUGUAAUGACAAAAGUAUCCAUAUUCUUGACAGUUAUUUUAUAUUAUAAAAAAAAGAUGAAGGAACUUUGACUGAUCUAAUAUGCUCAACGAAAGUGAAUCAAAUGUUAUAAAAUUACUUACACCACUAAUUGAAAACCUGUCUGAUAUUCAAUUAUUAUUUAUUAUUAUAUAAUUAUAUAAUAAUAAAUAAAAUGGUUGAUGUUAUGUAUUGGAAAUGAGCAUACGAUAAAUCAUAUAACCAUUAGUAAUAUAAUUUGAGAGCUAAGUUAGAUAUUUACGUAUUUAUGAUAAAACAGAAUAAACCCUAUAAAUUAUUAUUAUUAAUAAUAAAAAAUAAUAAUAAUACCAAUAUAUAUAUUAUUUAAUUUAUUAUUAUUAUAUUAAUAAAAUUUAAUAUAUAUUAUAAAUAAUUAUUGGAUUAAGAAAUAUAAUAUUUUAUAGAAAUUUUCUUUAUAUUUAGAGGGUAAAAGAUUGUAUAAAAAGCUAAUGCCAUAUUGUAAUGAUAUGGAUAAGAAUUAUUAUUCUAAAGAUGAAAAUCUGCUAACUUAUACUAUAGGUGAUAUGCCUAUCUUUAUUUAUAUAUAUAUUAUUAUUAUUAAUAAUAAAAAAAAAUUAAAAAAGAUAGGAGGUUUAUAUAUAACUGAUAAAUAUUUAUUAUAUUAUUUUUUUUAUAAUAAAUAUUAAAAGAUAUUGCGUGAGCCGUAUGCGAUGAAAGUCGCACGUACGGUUCUUACCGGGGGAAAACUUGUAAAGGUCUACCUAUCGGGAUACUAUGUAUUAUCA-3'

**Scheme S2.** RNA sequence of the full-length group IIB intron (942 nucleotides) with the shortened flanking exons ex^41^-*Sc*.ai5γ-ex^14^.

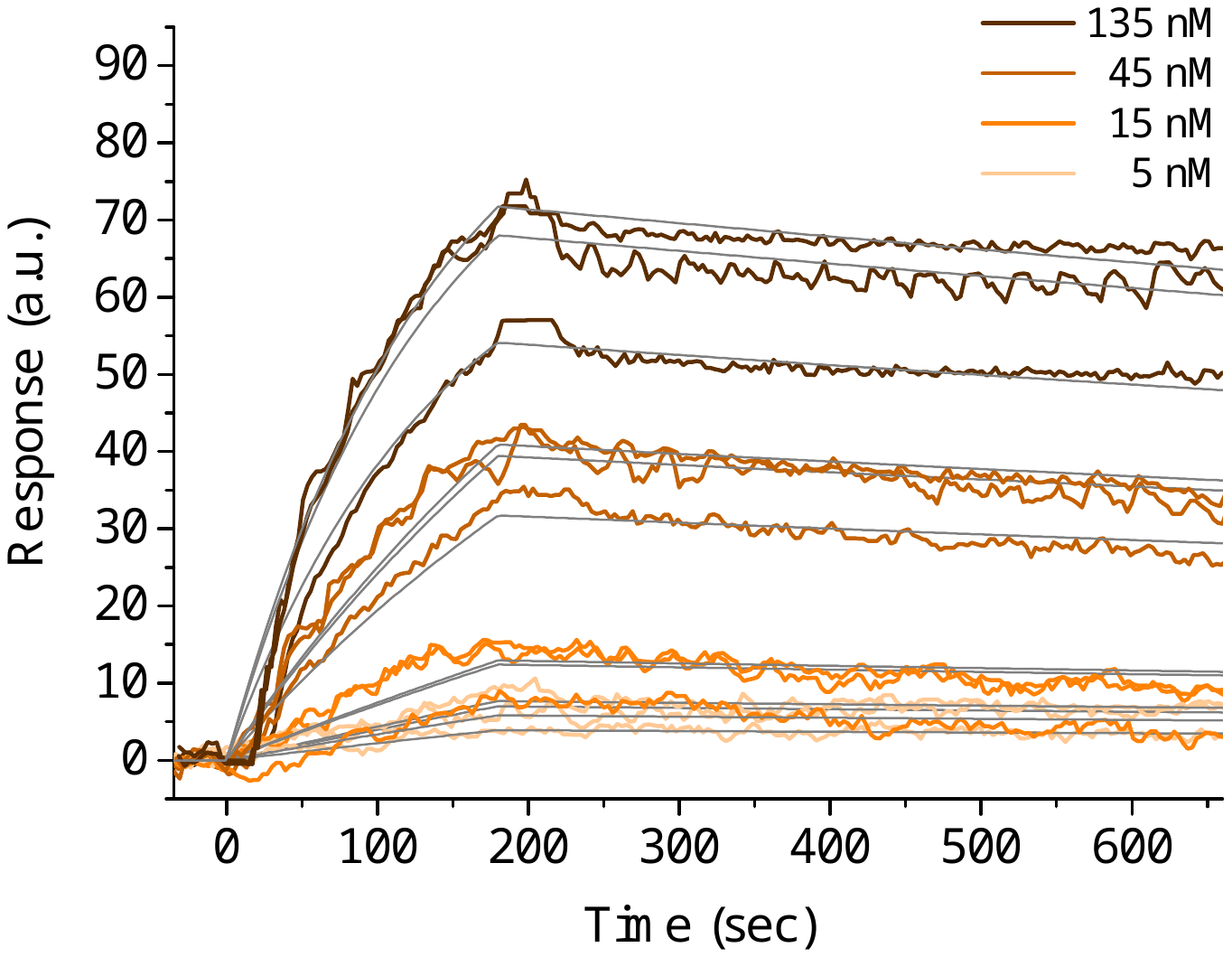

Fig. S2. SPRi measurements of the group IIB intron ex^41^-*Sc*.ai5γ-ex^14^ with a concentration series of Mss116 (5-135 nM, dilution factor = 3) in the absence of ATP in the running buffer. The calculated *K*_D_ = 5.40 ± 0.06 nM, with the fits shown in grey. The group IIB intron construct was immobilized via domain IV and a hybridized biotin-carrying DNA oligonucleotide.

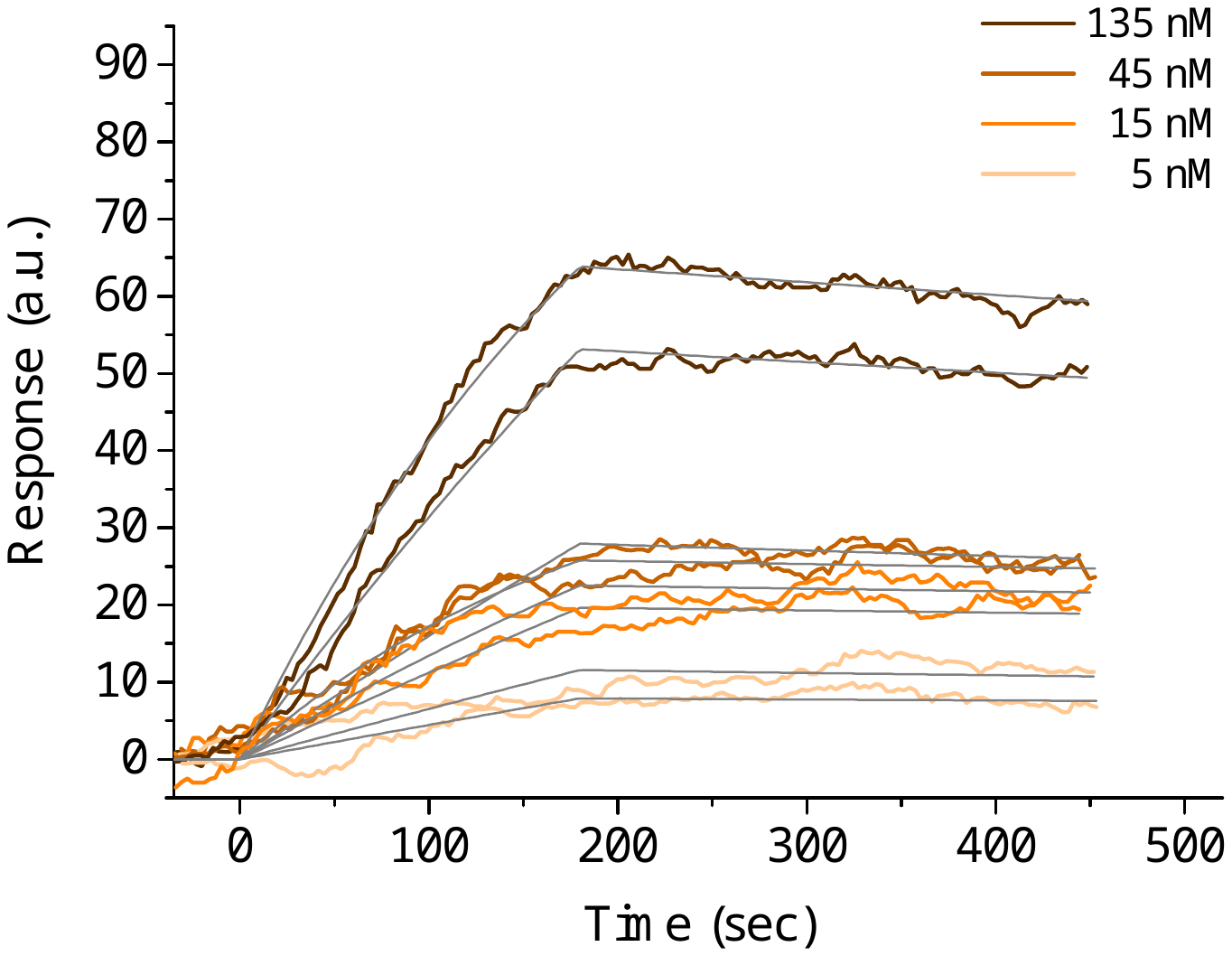

Fig. S3. SPRi measurements of the group IIB intron ex^41^-*Sc*.ai5γ-ex^14^ in the presence of ATP in the running buffer and a concentration series of Mss116 (5-135 nM, dilution factor = 3). The calculated *K*_D_ = 9.1 ± 0.3 nM, and the fits are shown in grey. The group IIB intron construct was immobilized at 5'exon position and a hybridized biotin-carrying DNA oligonucleotide.

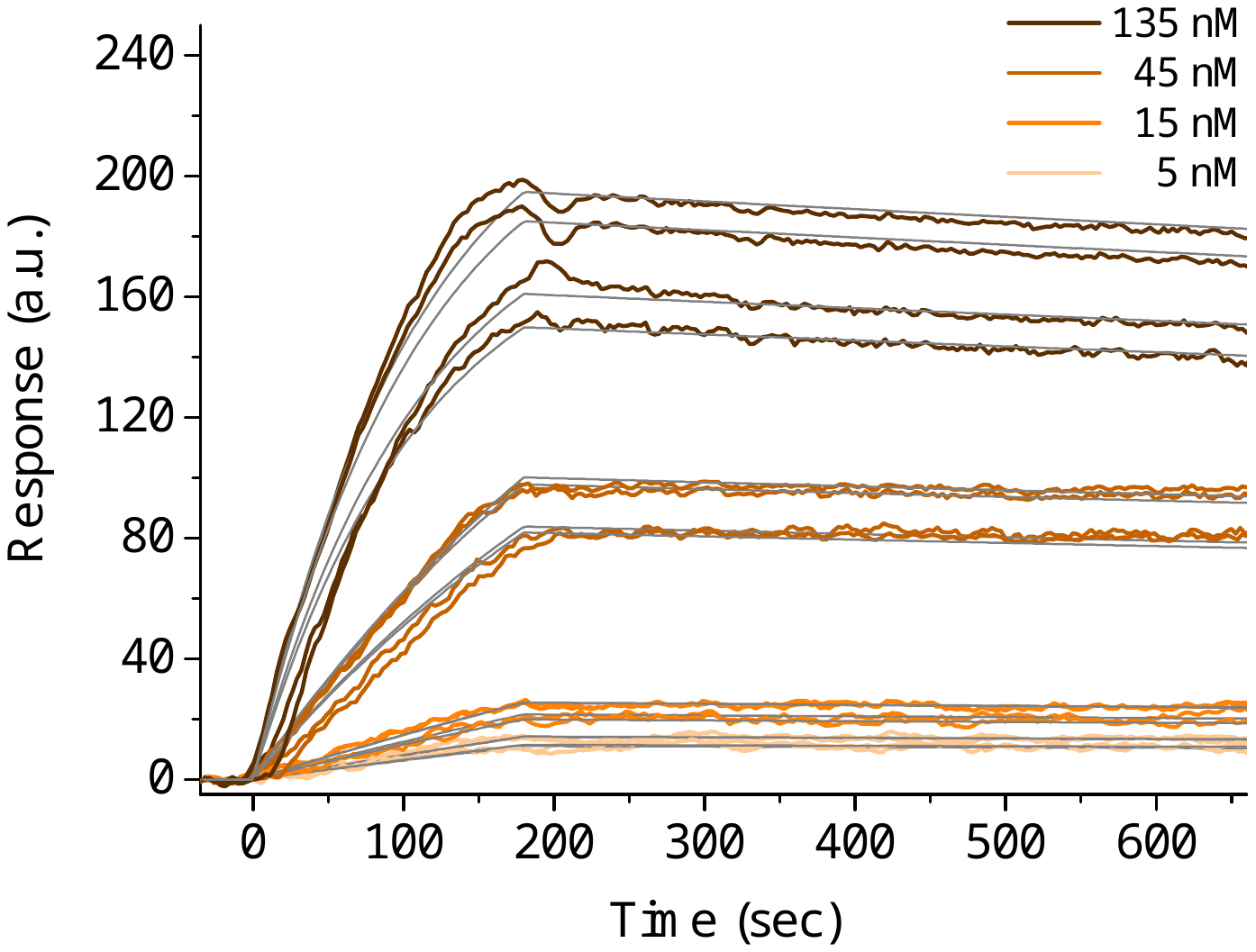

Fig. S4. SPRi measurements of the group IIB intron ex^41^-*Sc*.ai5γ-ex^14^ in the presence of ATP in the running buffer and a concentration series of Mss116 (5-135 nM, dilution factor = 3). The calculated *K*_D_ = 2.07 ± 0.02 nM, and the fits are shown in grey. The group IIB intron construct was immobilized via domain IV and a hybridized amine-carrying DNA oligonucleotide.

Supplementary Table S2. Overview of the peptide mass fingerprint analysis of on-plate digestion of bovine serum albumin (BSA) using different concentrations of reducing agents TCEP and DTT (reduction time 1 h). No alkylation step was performed.

| n(BSA) | c(TCEP) | Sequence coverage | Intensity coverage |
| --- | --- | --- | --- |
| 1 pmol  100 fmol  10 fmol | 50 mM | 0.0 %  0.0 %  0.0 % | 0.0 %  0.0 %  0.0 % |
| 1 pmol  100 fmol  10 fmol | 10 mM | 66.9 %  36.9 %  0.0 % | 74.8 %  23.0 %  0.0 % |
| 1 pmol  100 fmol  10 fmol | 5 mM | 51.2 %  38.9 %  0.0 % | 49.1 %  26.5 %  0.0 % |
| 1 pmol  100 fmol  10 fmol | 1 mM | 57.8 %  23.7 %  7.9 % | 79.2 %  46.5 %  11.6 % |
| 1 pmol  100 fmol  10 fmol | 0.5 mM | 72.7 %  62.1 %  42.7 % | 93.9 %  93.7 %  95.7 % |
| 1 pmol  100 fmol  10 fmol | 0.1 mM | 18.6 %  27.5 %  26.2 % | 65.3 %  97.2 %  98.0 % |
| 1 pmol  100 fmol  10 fmol | 0.01 mM | 29.8 %  28.7 %  29.0 % | 78.5 %  96.8 %  93.9 % |

| n(BSA) | c(DTT) | Sequence coverage | Intensity coverage |
| --- | --- | --- | --- |
| 1 pmol  100 fmol  10 fmol | 50 mM | 23.2 %  12.7 %  8.7 % | 82.3 %  78.6 %  74.4 % |
| 1 pmol  100 fmol  10 fmol | 10 mM | 35.7 %  35.3 %  0.0 % | 94.3 %  85.9 %  0.0 % |
| 1 pmol  100 fmol  10 fmol | 5 mM | 43.3 %  22.4 %  0.0 % | 74.8 %  91.4 %  0.0 % |
| 1 pmol  100 fmol  10 fmol | 1 mM | 40.7 %  18.1 %  16.5 % | 90.6 %  69.6 %  61.7 % |
| 1 pmol  100 fmol  10 fmol | 0.5 mM | 27.7 %  24.7 %  22.7 % | 94.4 %  90.8 %  91.8 % |
| 1 pmol  100 fmol  10 fmol | 0.1 mM | 22.7 %  23.1 %  22.7 % | 94.4 %  95.5 %  89.5 % |
| 1 pmol  100 fmol  10 fmol | 0.01 mM | 23.6 %  25.0 %  19.8 % | 90.7 %  85.2 %  87.2 % |

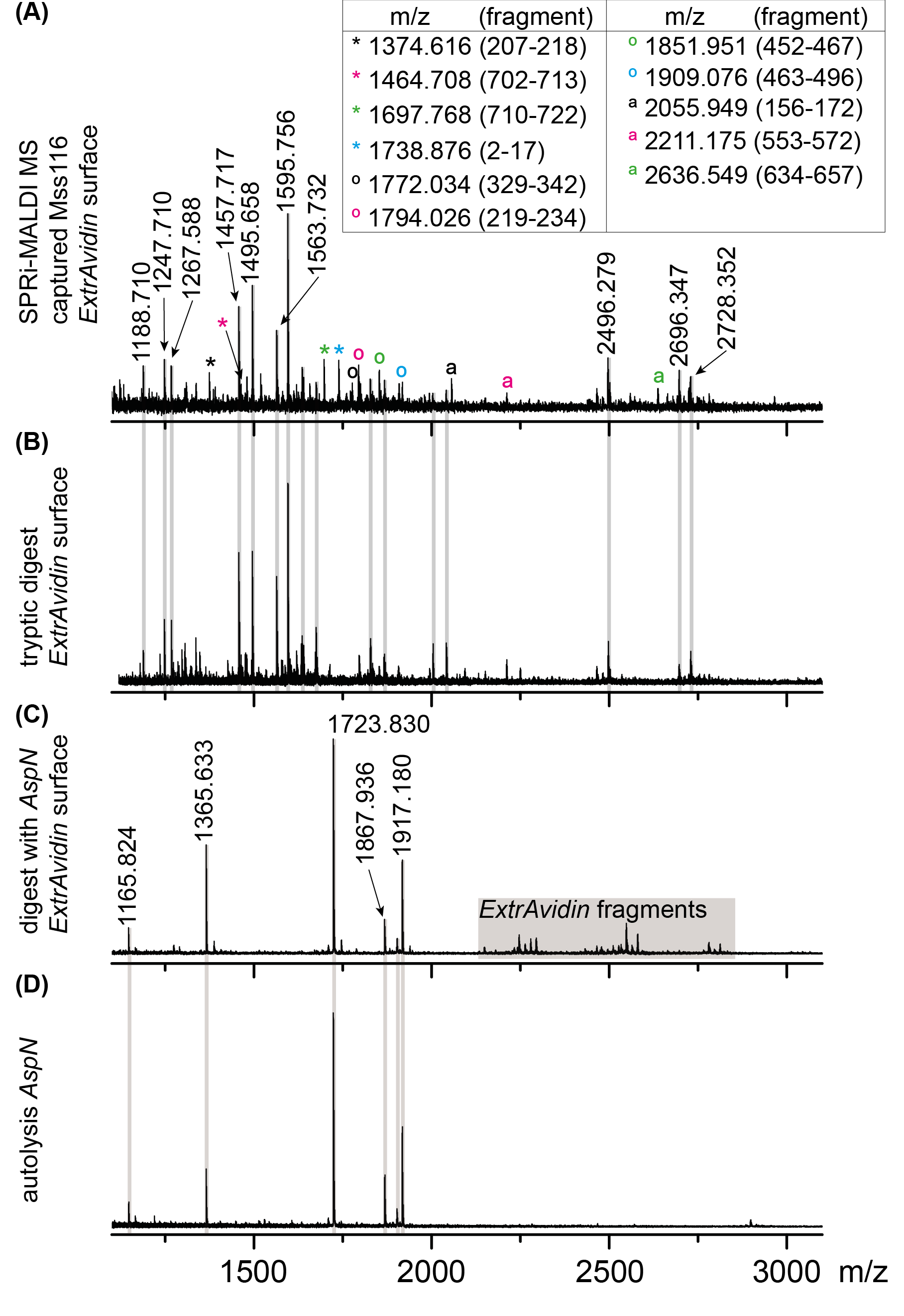

Fig. S5. SPRi-MALDI MS and MALDI MS control measurements. From top to bottom: SPRi-MALDI mass spectrum after injecting Mss116 while the group IIB intron ex^41^-*Sc*.ai5γ-ex^14^ was immobilized via DIV and a hybridized biotin-carrying DNA oligonucleotide (oMGG03) on an *ExtrAvidin* coated SPRi slide (A). Identified peptides from Mss116 after on-chip tryptic digestion are labeled with symbols in different colors and listed in the table on top (m/z value plus amino acid range in brackets). Unfortunately, the mass spectrum is dominated by peptides originating from the *ExtrAvidin* coating, which is also digested (B) (shown for comparison). Substituting the enzyme trypsin to AspN, *ExtrAvidin* fragments with higher m/z values (> 2100 Da) were expected and detected, but autolysis peptides of the enzyme are now dominating (C). The control autolysis of AspN (D) was performed on a commercial stainless steel MALDI target plate. Working with the same low concentration as used for trypsin or even lower while still being active did not minimize autolysis peptides of AspN.

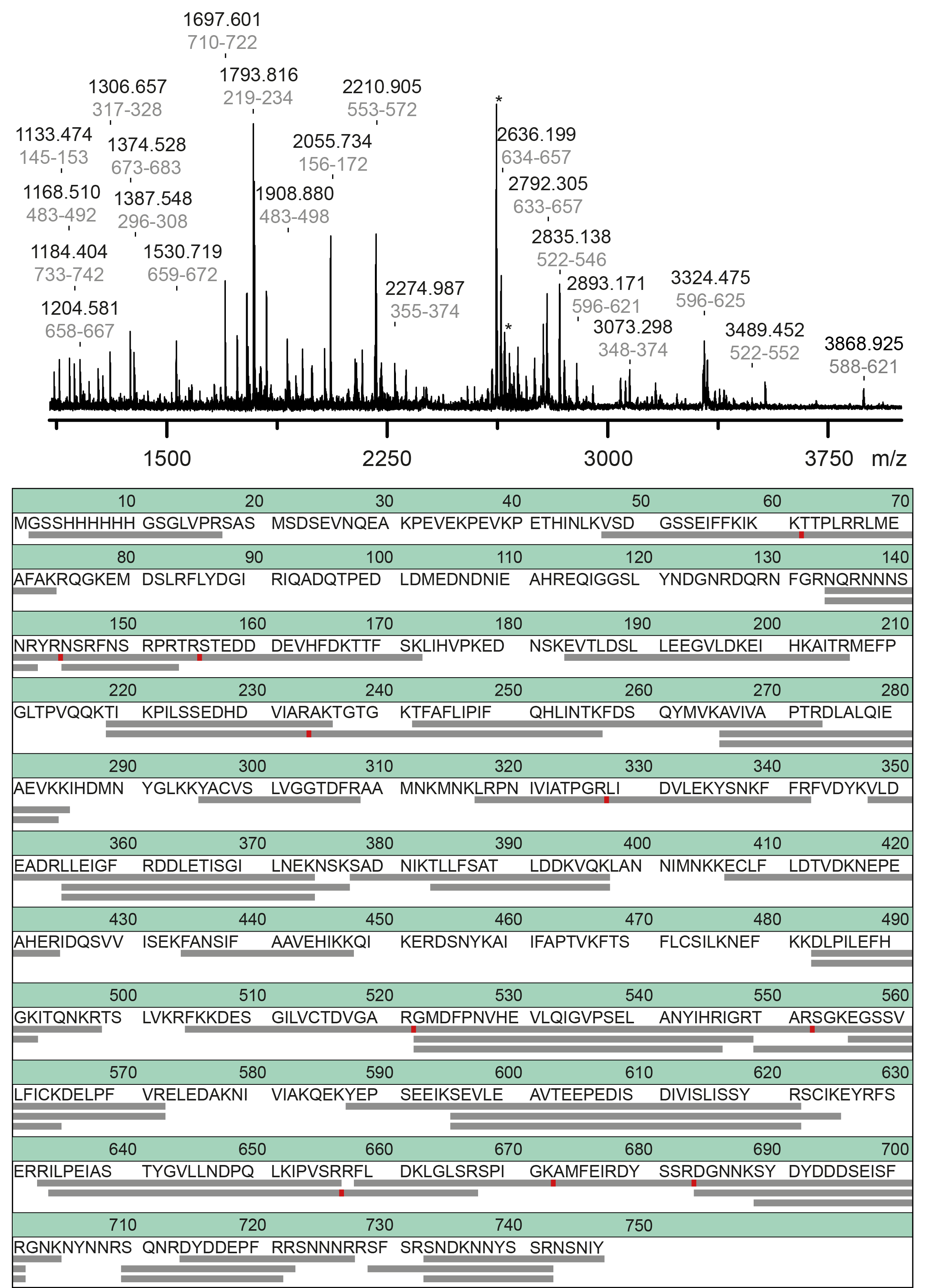

Fig. S6. On-chip MALDI mass spectrum after SPRi measurement from captured Mss116. Fragments of the on-chip proteolytic digested target protein are labeled within the spectrum with their m/z value and sequence range (in grey). All assigned ions show a higher signal-to-noise ratio than 1.5 and deviate between 60-190 ppm. For a better overview, all identified peptides are shown in the table below the spectrum – corresponding peptides and sequence ranges are marked with grey bars. The red point indicates a cleavage. Peaks marked with an asterisk origin from a contamination.

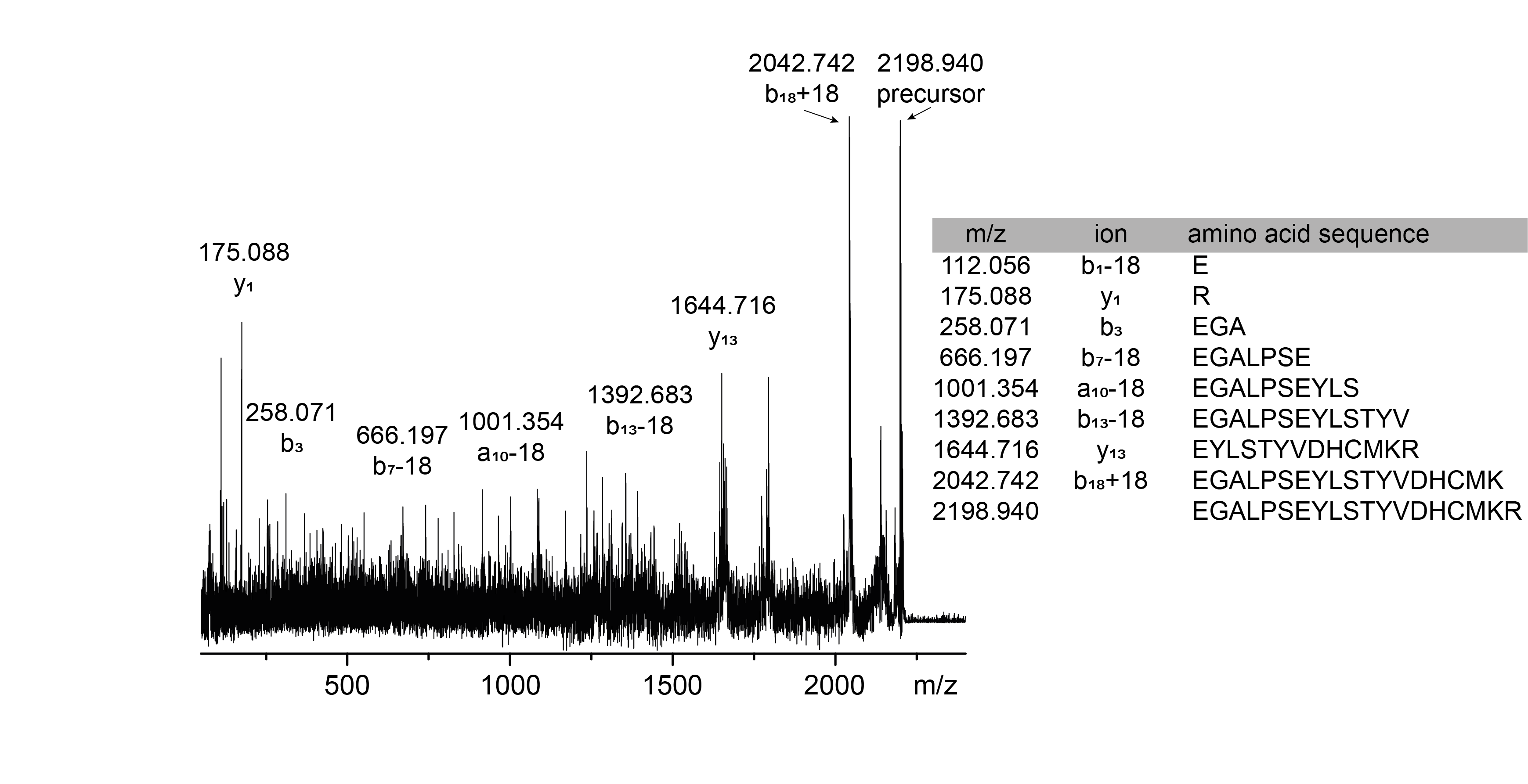

Fig. S7. SPRi-MALDI MS: On-chip MS/MS measurement of the peptide m/z 2198.940 (Cullin-3) detected after injecting a rough mitochondrial fraction lysate. A complete list of identified fragments (m/z values, cleavage site of the peptide ("ion") and positions of involved amino acids and amino acid sequence) is given in the table to the right to the spectrum.
